# Chromosome counting in the mouse zygote using low-invasive super-resolution live-cell imaging

**DOI:** 10.1101/2021.03.14.435348

**Authors:** Yu Hatano, Daisuke Mashiko, Mikiko Tokoro, Tatsuma Yao, Kazuo Yamagata

## Abstract

In preimplantation embryos, an abnormal chromosome number causes developmental failure and a reduction in the pregnancy rate. Conventional chromosome testing methods requiring biopsy reduce the risk of associated genetic diseases; nevertheless, the reduction in cell number also reduces the pregnancy rate. Therefore, we attempted to count the chromosomes in mouse (Slc:ICR) embryos using super-resolution live-cell imaging as a new method of chromosome counting that does not reduce the cell number or viability. We counted the forty chromosomes at the first mitosis by injecting embryos with histone H2B-mCherry mRNA under conditions by which pups could be obtained; however, the results were often an underestimation of chromosome number and varied by embryo and time point. Therefore, we developed a method to count the chromosomes via CRISPR/dCas-mediated live-cell fluorescence *in situ* hybridization targeting the sequence of the centromere region, enabling us to count the chromosomes more accurately in mouse embryos. The methodology presented here may provide useful information for assisted reproductive technologies, such as those used in livestock animals/humans, as a technique for assessing the chromosomal integrity of embryos prior to transfer.

**Impact Statement:** Low-toxic super-resolution observation enables chromosome counting in preimplantation embryos without cell collection.

## Introduction

In preimplantation embryos, the number of chromosomes is correlated with the pregnancy rate in assisted reproductive technology (ART) and animal breeding (***Magli et al., 2000, Sandalinas et al., 2001, Rubio et al., 2007, Mantikou., 2012, Yao et al., 2018***). In addition, even if embryos with abnormal chromosome numbers achieve full-term development, they may exhibit severe genetic disease (e.g., trisomy 21) (***Lejeune et al., 1959***). Therefore, evaluating chromosome numbers prior to embryo transplantation is important to mitigate these risks. Conventional chromosomal testing methods such as collecting and disrupting cells, subsequent spreading of the chromosomes, or scanning the genome using microarrays or next-generation sequencing can reduce the risk of genetic disease (Hens et al., 2013). Meanwhile, an associated risk of decreased pregnancy rate consequent to cell harvesting from chromosomally normal embryos has been reported **(*Mastenbroek et al., 2007*)**. As approximately 90% of the causes of trisomy 21 are derived from meiosis (***Antonarakis et al., 1991, Freeman et al., 2007***), determining the number of chromosomes immediately following fertilization may facilitate the application of strategies to reduce the risk of genetic disease by excluding embryos with abnormal chromosome numbers from transplantation. Nevertheless, the number of chromosomes in early-stage embryos cannot be evaluated using these techniques since they are conventionally performed using blastocysts, which contain a sufficient number of cells to support cell collection. Therefore, the development of technology that enables real-time testing of chromosomes immediately after fertilization without reducing the number of pre-implantation germ cells will represent a breakthrough that will bring new insights into ART.

Live-cell imaging constitutes a technology for observing the interior of cells in their native environment. Through low-toxic, long-term live-cell imaging of mouse early embryos using fluorescence microscopy (***Yamagata 2009a***), we previously demonstrated that the early division of preimplantation embryos affects subsequent ontogeny. For example, abnormal chromosome segregation during early division affects development up to the blastocyst stage (***Mashiko et al., 2020***), resulting in extremely low birth rates following transplantation of 2-cell stage embryos (***Yamagata et al., 2009b***). However, although obvious abnormalities leading to the formation of micronuclei could be detected, it was not possible to count the number of chromosomes owing to limited resolution (***Mashiko et al., 2020***).

In recent years, super-resolution microscopes based on various principles have been proposed and used for imaging cells (i.e., stimulated emission depletion microscopy (STED): ***Hell et al., 1994**; **Hell et al., 1995***; photoactivated localization microscopy (PALM): ***Betzig et al., 2006***; fluorescence photoactivation localization microscopy (FPALM): ***Hess et al., 2006***; stochastic optical reconstruction microscopy (STORM): ***Rust et al., 2006***; structured illumination microscopy (SIM): ***Gustafsson et al., 2000***; and Airyscan: ***Huff, 2015***). Nevertheless, long-term observation using a super-resolution microscope may be considered as highly invasive to observed cells. In particular, during the observation of preimplantation embryos, acquisition of three-dimensional images and repeated irradiation by the laser for time-lapse imaging causes embryo damage. Thus, fluorescence observation under conditions that allow the development of embryos to term without prior arrest is important to ensure that the observed phenomenon is not an artifact caused by dying cells. To this end, we attempted to identify appropriate long-term live-cell imaging conditions using a disk confocal type fluorescence microscope to observe early embryogenesis and predict prognosis (***Yamagata et al., 2009a***). In general, confocal microscopy scans a sample at a single plane, thus requiring substantial time to obtain an image. In addition, application of high-intensity laser energy may readily discolor the sample or have cytotoxic effects. In comparison, spinning disk confocal microscopy can overcome these limitations by scanning multiple points through rotation of the disk and is therefore suitable for live-cell imaging of early embryos. Moreover, super-resolution has recently been realized using a disk confocal, optical photon reassignment microscopy (OPRA)-type microscope by optically reducing individual focal points projected on pinholes using microlenses (super-resolution via optical re-assignment (SoRa): ***Azuma and Kei, 2015***).

The objective of this study is therefore to develop a technology capable of performing chromosome counts in preimplantation embryos without requiring cell collection to identify embryos with aberrant chromosome numbers. Using mice, which have successfully been used as model animals for the study of mammalian embryos and the establishment of associated techniques, we applied the SoRa system to count chromosomes of early-stage embryos under conditions that allowed the embryo to maintain full-term developmental potential following the super-resolution observation.

## Results

### Construction of the SoRa live-cell imaging system and optimization of imaging conditions with minimal phototoxicity

We constructed a disk confocal type super-resolution live-cell imaging system to observe mouse fertilized eggs (***Fig. 1***). As the system has a switching mechanism between the spinning disk with (SoRa disk) and without (Normal disk) microlenses, we can observe the same specimen under these two different modes. To cover the broad viewing field of the system, a scientific complementary metal–oxide–semiconductor (sCMOS) image sensor was used and denoise algorithms (***Boulanger et al., 2009***) were activated at every acquisition to compensate for the lower sensitivity compared to electron multiplying charge-coupled devices (EMCCD) sensor. For observation of the first mitosis (approximately 17 h from the pronuclear stage to the end of the first mitosis), an incubator and gas chamber were set up on the stage for culturing embryos. A constant room temperature was maintained at 30 °C so that the embryos would not be affected by outside air.

**Figure 1.**
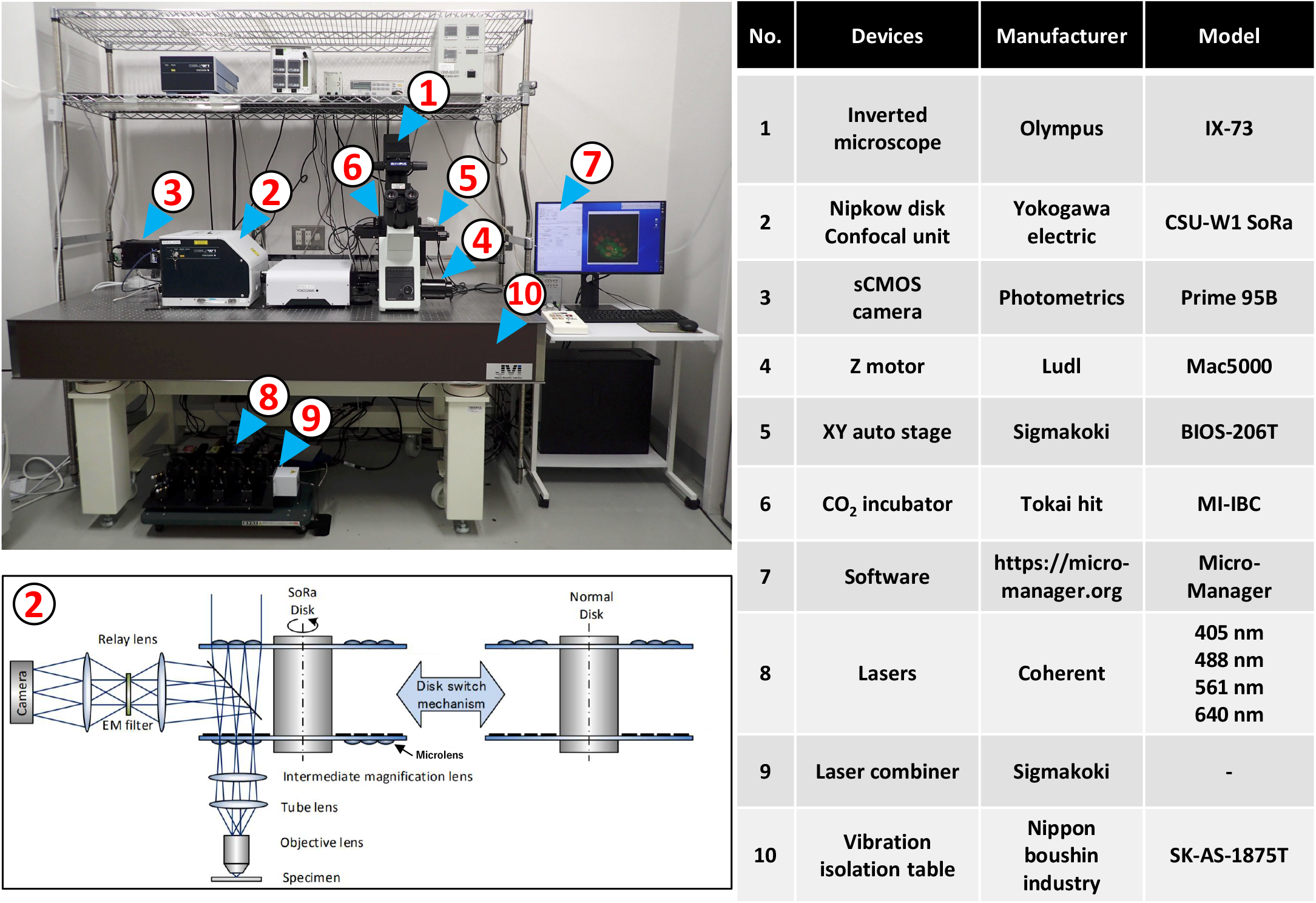
SoRa live-cell imaging system. Photograph (left) and list (right) of the equipments for imaging. A conventional inverted microscope was attached to a Nipkow disk confocal unit, sCMOS camera, Z-motor, and XY auto stage. All devices were controlled using a micro-manager. Embryos were cultured in a CO_2_ incubator at this stage. The Nipkow disk confocal unit can switch between a conventional unit without microlenses and a unit with microlenses (lower left).

To evaluate the phototoxicity resulting from observation using the constructed SoRa live-cell imaging system, we focused on the decay of the brightness (fading) of H2B-mCherry during the observation of early embryos. We injected mRNA encoding histone H2B-mCherry into fertilized eggs. One plane of the z-axis of the nucleus of the mouse 2-cell stage embryo was observed for 100 s continuously in streaming mode. Despite observation under the same conditions, such as excitation laser power, exposure time, and camera sensitivity, a decrease in signal was observed with SoRa compared to that observed using the conventional disk confocal microscopy system W1 (time constant 82.4 vs 137 ms, respectively) (***Fig. 2A, B***). This result suggested that the SoRa system requires more stringent conditions to suppress the phototoxicity than the conventional W1 system, which likely arises because the SoRa system has a higher light density (power of laser per unit area) than the conventional W1 system.

**Figure 2.**
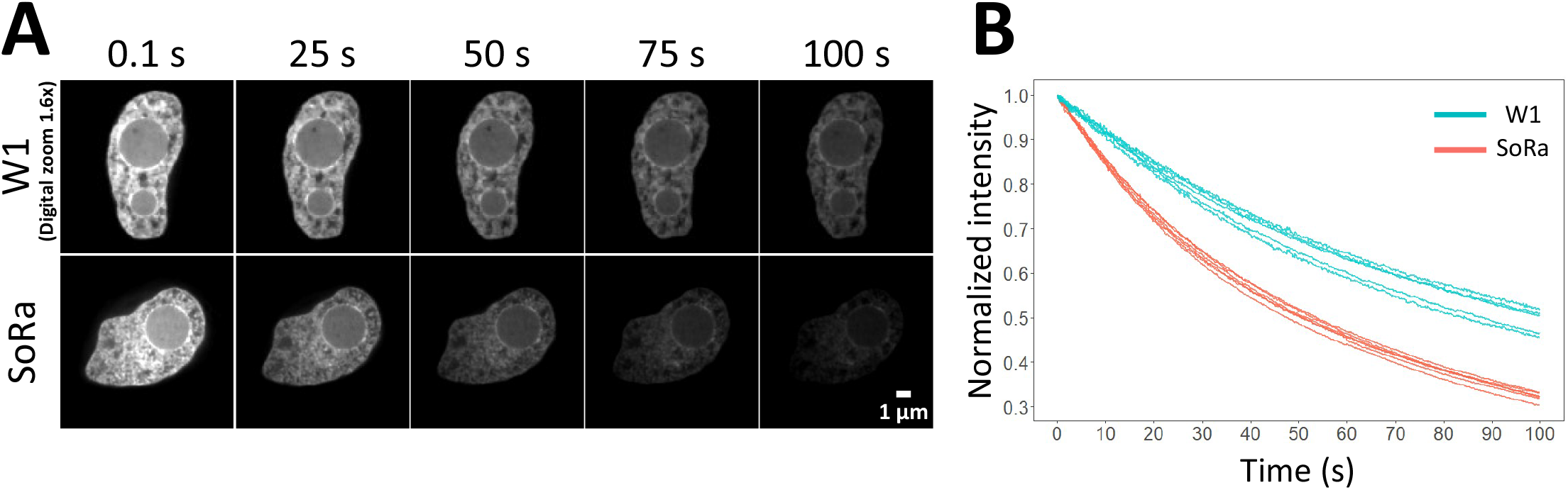
Assessment of the phototoxicity of super-resolution imaging. (**A**) Photographs of 2-cell nuclei obtained using the W1 (upper panels) and SoRa (below panels) systems. bar: 1 μm. (**B**) Graph of the brightness change.

We next searched for conditions that did not affect embryogenesis following observation with a super-resolution microscope (***Table. 1***). We examined the wavelength, laser power, and time intervals. At an excitation wavelength of 561 nm, laser power of 0.1 mW (emitted from tip of objective), and 5 min intervals, the number of embryos that reached the 2-cell stage was 19/20 (95%) and 14/20 (70%) reached the blastocyst stage. At an excitation wavelength of 561 nm, laser power of 0.1 mW, and 10 min intervals, 18/20 (90%), embryos reached 2 cells and 16/20 (80%) reached the blastocyst stage. The blastocyst arrival rate under these conditions did not differ from that in the non-observed group (52/65 (80%), *P* = 0.89, *P* = 1, prop-test). Laser power of 0.2 mW affected the growth up to the blastocyst stage more than 0.1 mW (0.2 mW, 5 min: 0/10 (0%); 0.2 mW, 10 min: 3/19 (15.8%), prop-test *P* = 0.016, 0.013, respectively). These results suggested that a laser power of 0.1 mW is suitable for an excitation wavelength of 561 nm. However, at an excitation wavelength of 488 nm and laser power of 0.1 mW, no embryos reached the blastocyst stage regardless of the time interval, indicating that an excitation wavelength of 488 nm is more toxic. We therefore utilized the conditions supportive of normal embryo development following super-resolution observation using a wavelength of 561 nm in subsequent experiments.

**Table 1.** Developmental capacity of the preimplantation embryo following super-resolution imaging. From left to right column: injected mRNA, excitation wavelength of the laser (nm), optical intensity (mW), time interval (min), imaging period (h), number of embryos examined, number of 2-cell embryos, and number of blastocysts.

| Injected mRNA | Optical wavelength (nm) | Optical intensity (mW) | Time intervals (min) | Imaging period (hour) | No. of embryos examined | No. (%) of 2-cell embryos | No. (%) of blastocyst |
| --- | --- | --- | --- | --- | --- | --- | --- |
| Histone H2B-mCherry | - | - | - | - | 65 | 63 (96.9) | 52 (80) |
|  | 561 | 0.1 | 5 | 17 | 20 | 19 (95) | 14 (70) |
|  | 561 | 0.1 | 10 | 17 | 20 | 18 (90) | 16 (80) |
|  | 561 | 0.2 | 5 | 17 | 10 | 4 (40) | 0 (0) |
|  | 561 | 0.2 | 10 | 17 | 19 | 15 (78.9) | 3 (15.8) |
| Histone H2B-EGFP | - | - | - | - | 7 | 7 (100) | 7 (100) |
|  | 488 | 0.1 | 5 | 17 | 5 | 1 (20) | 0 (0) |
|  | 488 | 0.1 | 10 | 17 | 5 | 5 (100) | 0 (0) |

To examine whether super-resolution live-cell imaging allowed subsequent full-term embryo development, 2-cell embryos observed at an excitation wavelength of 561 nm, laser power of 0.1 mW, and interval of 5 or 10 min were transplanted into the oviduct of pseudopregnant mice. Notably, pups were obtained under both intervals (5 min (1/10): *P* = 0.57, 10 min (7/15): *P* = 0.66, respectively vs. imaging (-) prop-test) (***Table. 2; Fig S1***). We therefore adopted a 10 min interval as the preferred observation condition.

**Table 2.** Full-term development of transferred embryos following super-resolution imaging. From left to right column: injected mRNA, excitation wavelength of the laser (nm), optical intensity (mW), time interval (min), imaging period (h), number of embryos examined, number of 2-cell stage embryos, number of recipients (transferred mice), number of transferred embryos, and number of pups. See also Supplementary figure 1.

| Injected mRNA | Optical wavelength (nm) | Optical intensity (mW) | Time intervals (min) | Imaging period (hour) | No. of embryos examined | No. (%) of 2-cell embryos | No. of recipients | No. (%) of Pups |
| --- | --- | --- | --- | --- | --- | --- | --- | --- |
| Histone H2B-mCherry | - | - | - | - | 27 | 26 (96.3) | 6 | 8 (29.6) |
|  | 561 | 0.1 | 5 | 17 | 13 | 10 (76.9) | 3 | 1 (7.7) |
|  | 561 | 0.1 | 10 | 17 | 15 | 15 (100) | 3 | 7 (46.7) |

Under the condition that the pups could be obtained (i.e., excitation wavelength 561 nm, laser power 0.1 mW, 10 min interval), time-lapse observation was then performed for 17 h until the completion of the first mitosis (***Fig. 3A, B; Movie. 1***).

**Figure 3.**
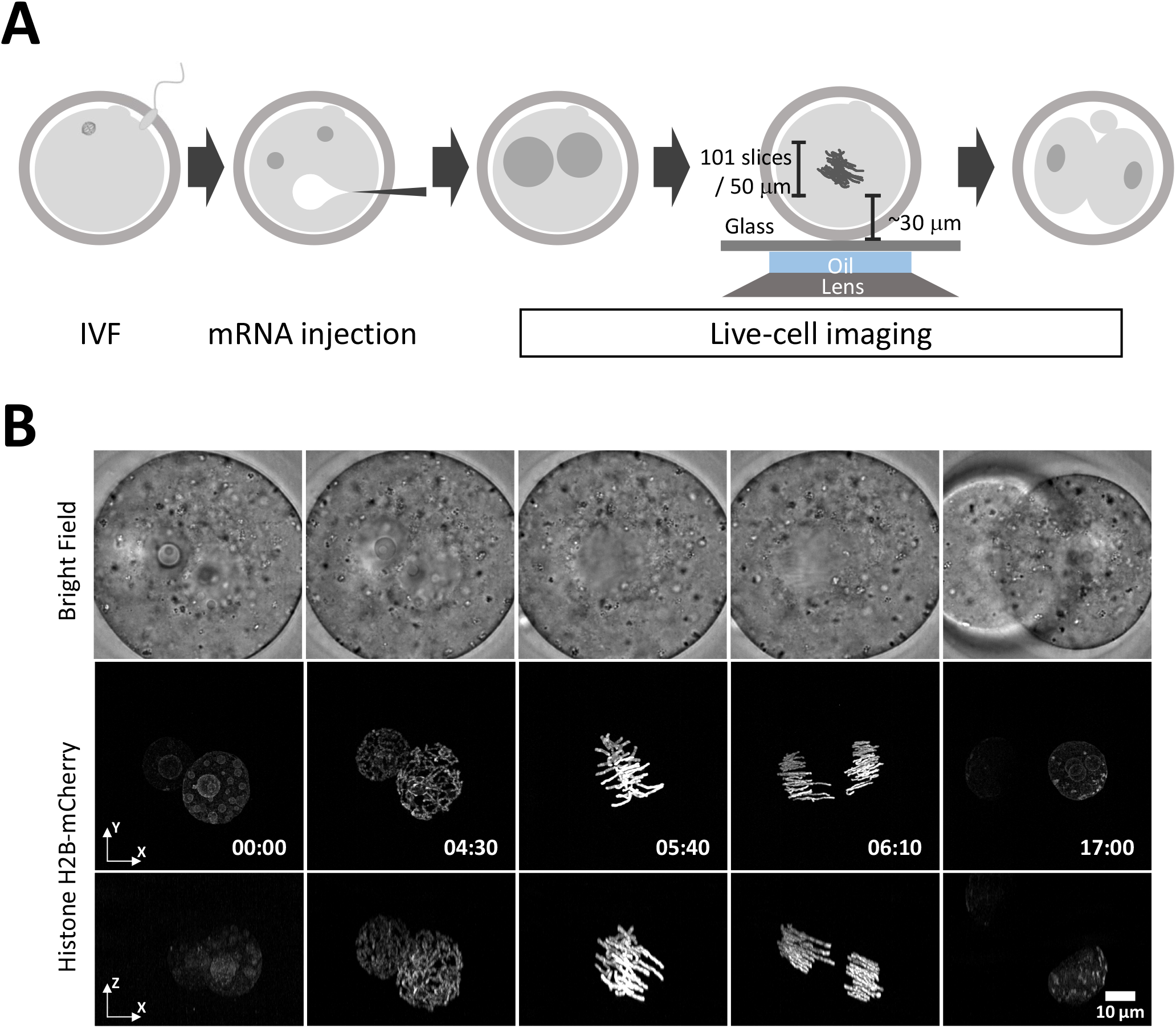
Super-resolution live-cell imaging of the first mitosis using chromosome counting by histone H2B-mCherry mRNA was the SoRa system. (**A**) Schematic diagram of super-resolution imaging of the first mitosis. (**B**) Super-resolution imaging of the first mitosis in mouse embryos. The upper panels show the bright-field images. Middle panels show the histone H2B-mCherry images of the x–y plane, and the panels below show that of the x–z plane. See also Supplementary movie 1.

### SoRa system allows observation of the grooves between the sister chromatids

Subsequently, the resolution of the SoRa system was compared with that of the W1 system using movies captured by each system. Chromosomes could be seen as two separating sister chromatids by observation using SoRa but not the conventional W1 system. Furthermore, by performing deconvolution processing on the images acquired with SoRa, the grooves of sister chromosomes could be separated more clearly (peak-to-peak length: 618.7 ± 137.5 nm) (***Fig. 4***). When interphase nuclei were observed, the contrast of the chromatin in the nuclei could be observed in more detail by imaging using the SoRa system than when using the conventional W1 system (***Fig. 4***).

**Figure 4.**
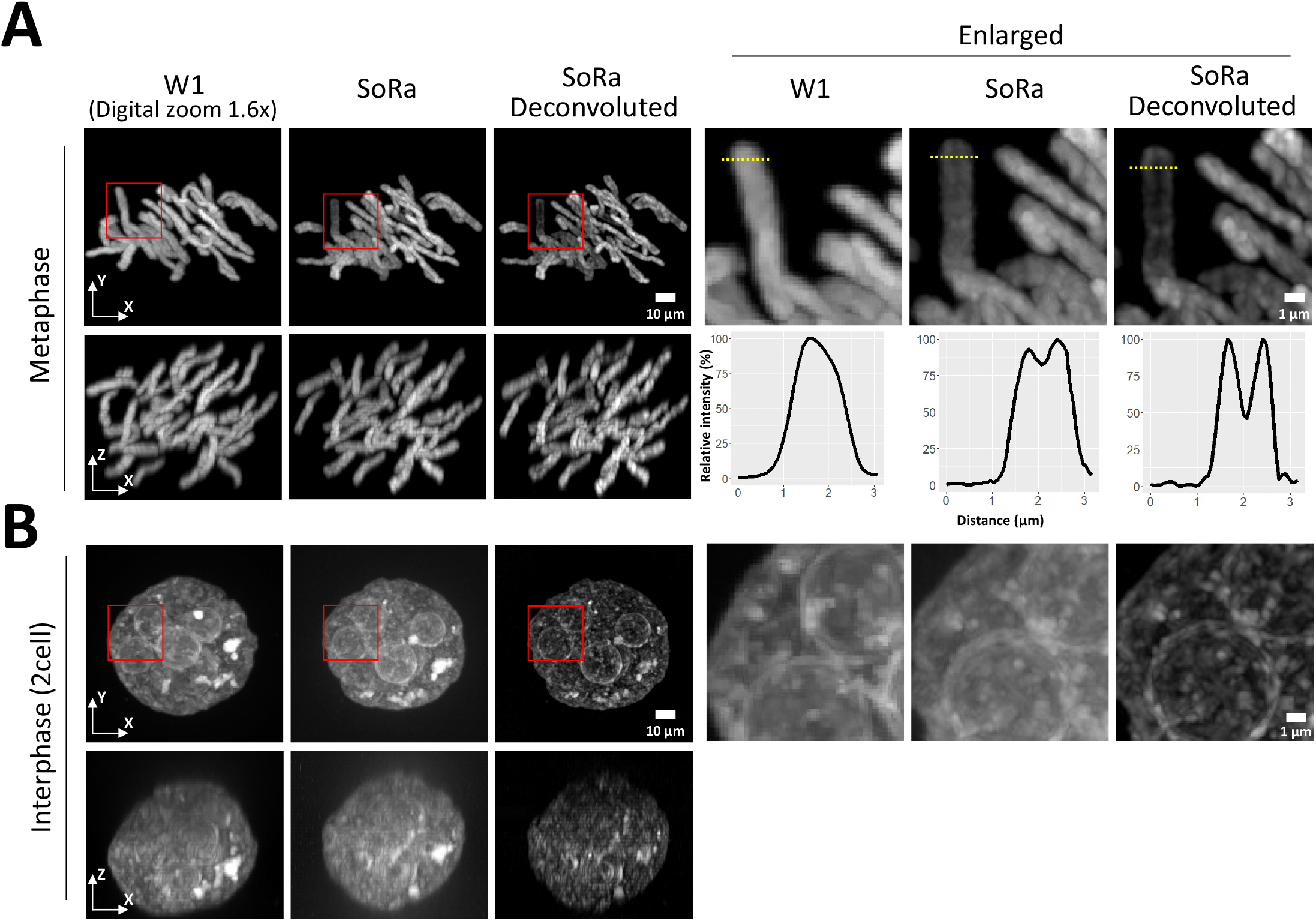
The SoRa system enables higher resolution imaging than the conventional W1 system. (**A**) Left six panels show images of the metaphase of an embryo. From left to right, the image was obtained using W1 (x–y and x–z), SoRa (x–y and x–z), and SoRa (x–y and x–z; deconvoluted). The right three panels show enlargements of the rectangular area (red). The right three graphs show the relative intensity on the yellow dashed line. (**B**) The left six panels show images of the interphase of a 2-cell embryo. The right three panels show enlargements of the rectangular area (red).

### Chromosome segmentation and auto-counting

For counting the number of chromosomes, we attempted the segmentation of M-phase chromosomes in living cells. Deconvoluted images were acquired using the Tikhonov regularization algorithm (***Sage et al., 2017***). Images were noise-processed using the Top Hat filter (***Legland et al., 2016***) and binarized with Otsu’s method (***Otsu, 1979***). As a result of automatically counting binarized objects (***Fig. 5A; Movie. 2***), we were able to count 40 objects (***Fig. 5B***). However, a risk existed of recognizing two objects as one when the chromosomes were close; consequently, the number of chromosomes detected were varied among the embryos and time points (***Fig. 5C***). Although the chromosomes were clearly separated compared to the results from conventional W1 system observation (***Fig. 5; Fig. S2***), the inconsistent counting of individual chromosomes represented a limitation of this technique.

**Figure 5.**
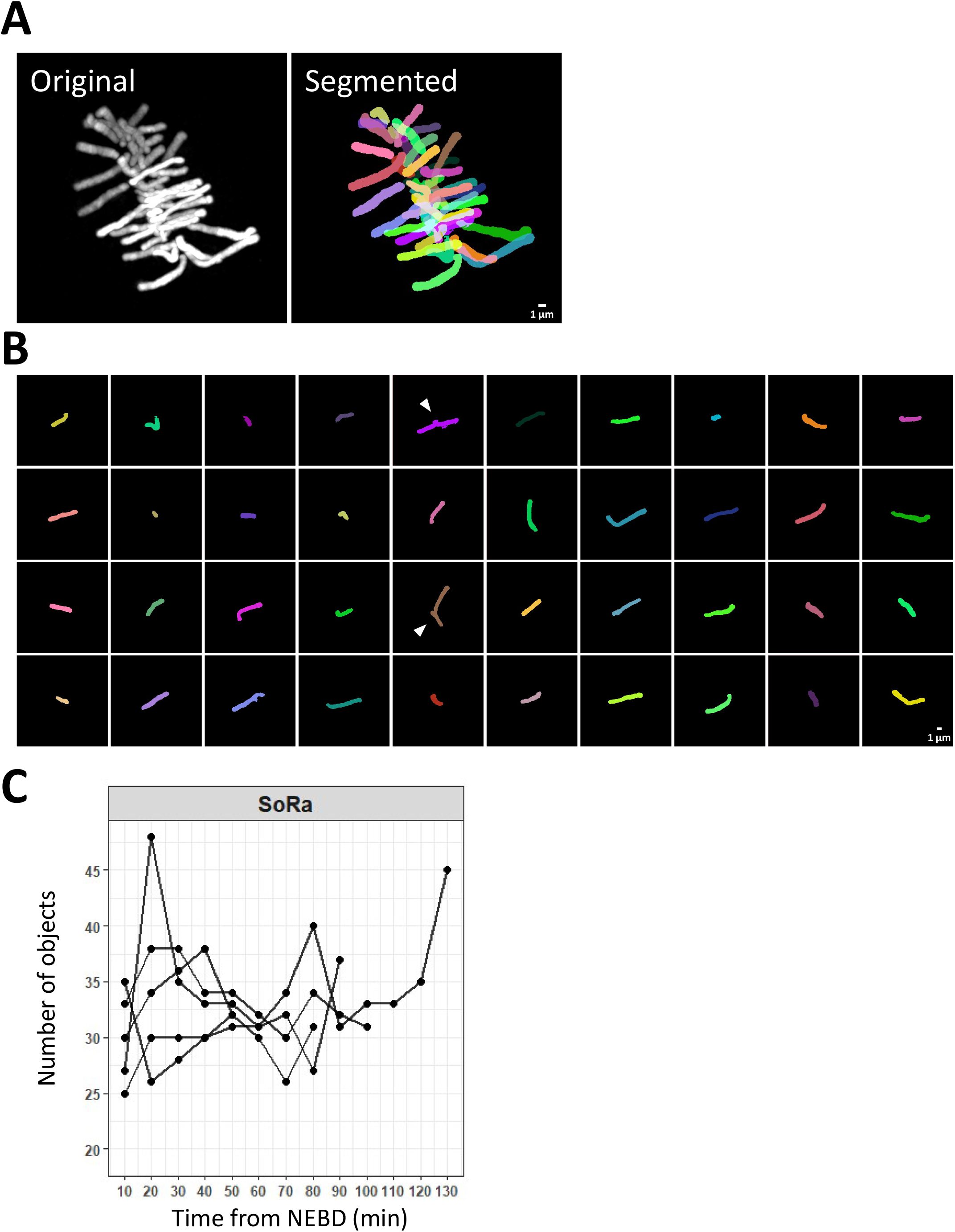
Chromosome counting using histone H2B-mCherry mRNA. (**A**) The left panel shows the histone H2B-mCherry signal. The right image shows the processed image. (**B**) Each panel shows the recognized object. The white arrowheads indicate a processed chromosome with a risk of miscount. (**C**) Relationship between the number of recognized objects and time from nuclear envelope breakdown (NEBD). See also Supplementary Movie 2 and Supplementary Figure 2.

### Chromosome counting by centromere labeling with live-fluorescence in situ hybridization (live-FISH) technology

To overcome the issue of miscounting at sites of overlapping chromosome signals, we sought to visualize the centromere, a more restricted DNA region in which the distal ends of all chromosomes are located (***Fig._6A***). Toward this end, CRISPR-mediated live imaging technology (***Wang et al., 2019***) was performed by targeting the minor satellite sequence, a repeat sequence constituting the centromere (***Fig. 6B***). First, we designed the crRNA based on the target sequence used in a previous report of ES cells (***Anton et al., 2014***). As counter staining, major satellite sequence locating pericentromere, a region on a chromosome that is larger and existing inward than the centromere (***Fig._6A***), was also labeled using TALE technology (***Miyanari et al, 2013***). As a result, the signals of both minor and major satellites were observed at the distal ends of metaphase and anaphase chromosomes (***Fig._6C***). Importantly, when compared to major satellites, the dot-like signals of minor satellites were more restricted, and overlapping satellites were not detected (***Fig. 6C***).

**Figure 6.**
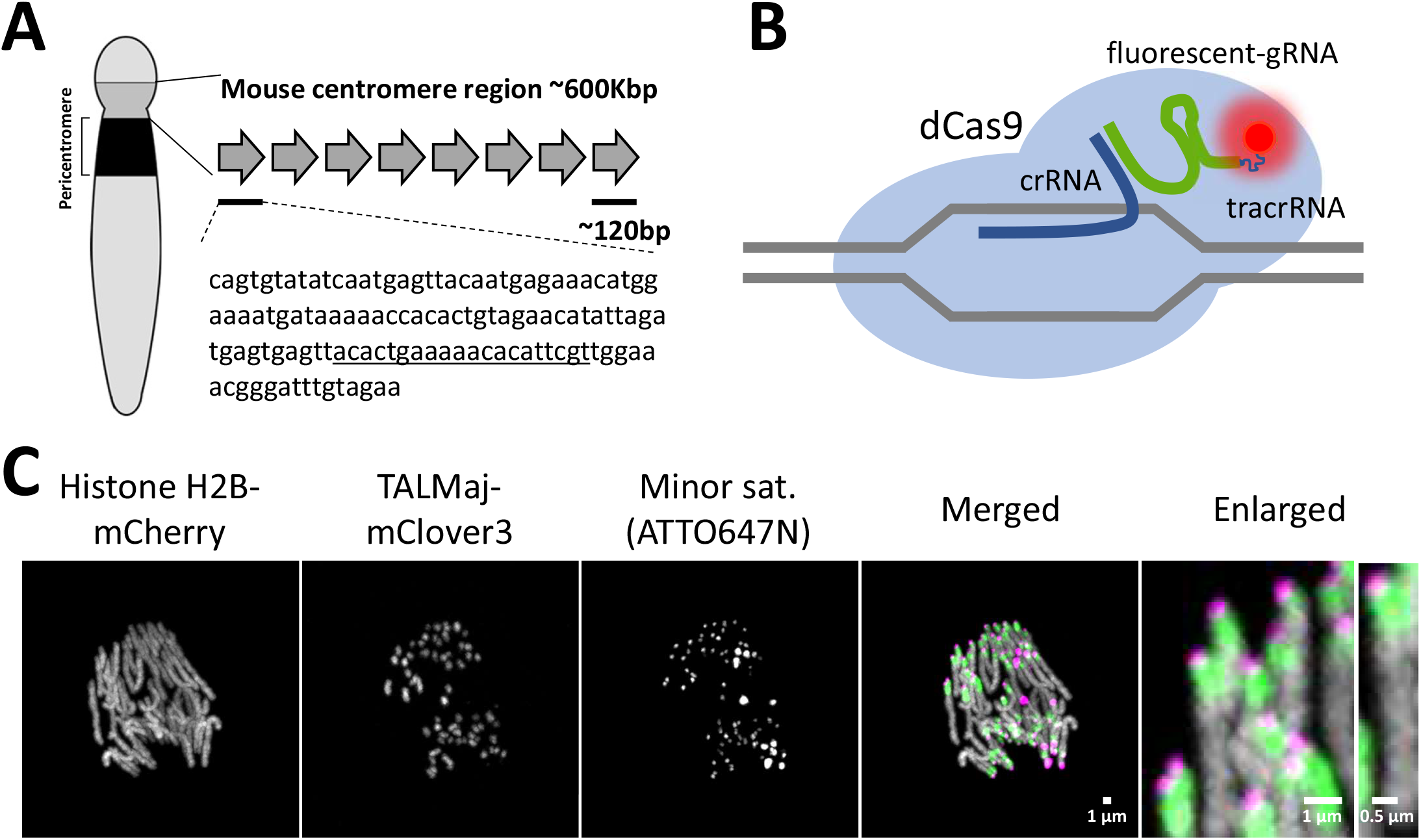
Chromosome counting by CRISPR-mediated live-FISH. (**A**) The left panel shows the illustrated chromosome; the grey and black areas show the mouse centromere and pericentromere regions, respectively. The below shows the repeated sequence of minor satellite; the underlined sequence shows the sequence targeted by gRNA. (**B**) Illustrated scheme of CRISPR-mediated live-cell imaging. The ATTO550 or 647N probe was added to tracrRNA. (**C**) Snapshot of Histone H2B-mCherry, TALMaj-mClover3, and minor satellite targeted gRNA using the SoRa system.

Finally, the number of chromosomes at metaphase were counted by combining centromeres labeled by live-FISH and the SoRa system. Consequently, it was possible to count 40 pairs of dot-shaped signals (***Fig. 7A, B; Movies. 3, 4***). Notably, the number of counted dot-like signals were 40 in all three embryos analyzed, which did not change with time (***Fig. 7C; Fig. S3***). In contrast, conventional W1 system did not consistently detect the dot-like signal (***Fig. 7; Fig. S4***). Moreover, through multicolor live-cell imaging using histone H2B-mCherry and ATTO647N-labeled gRNA for the minor satellite, we found that embryos would reach the blastocyst stage (3/6, ***Fig. 7; Fig. S5, Movie. 5***).

**Figure 7.**
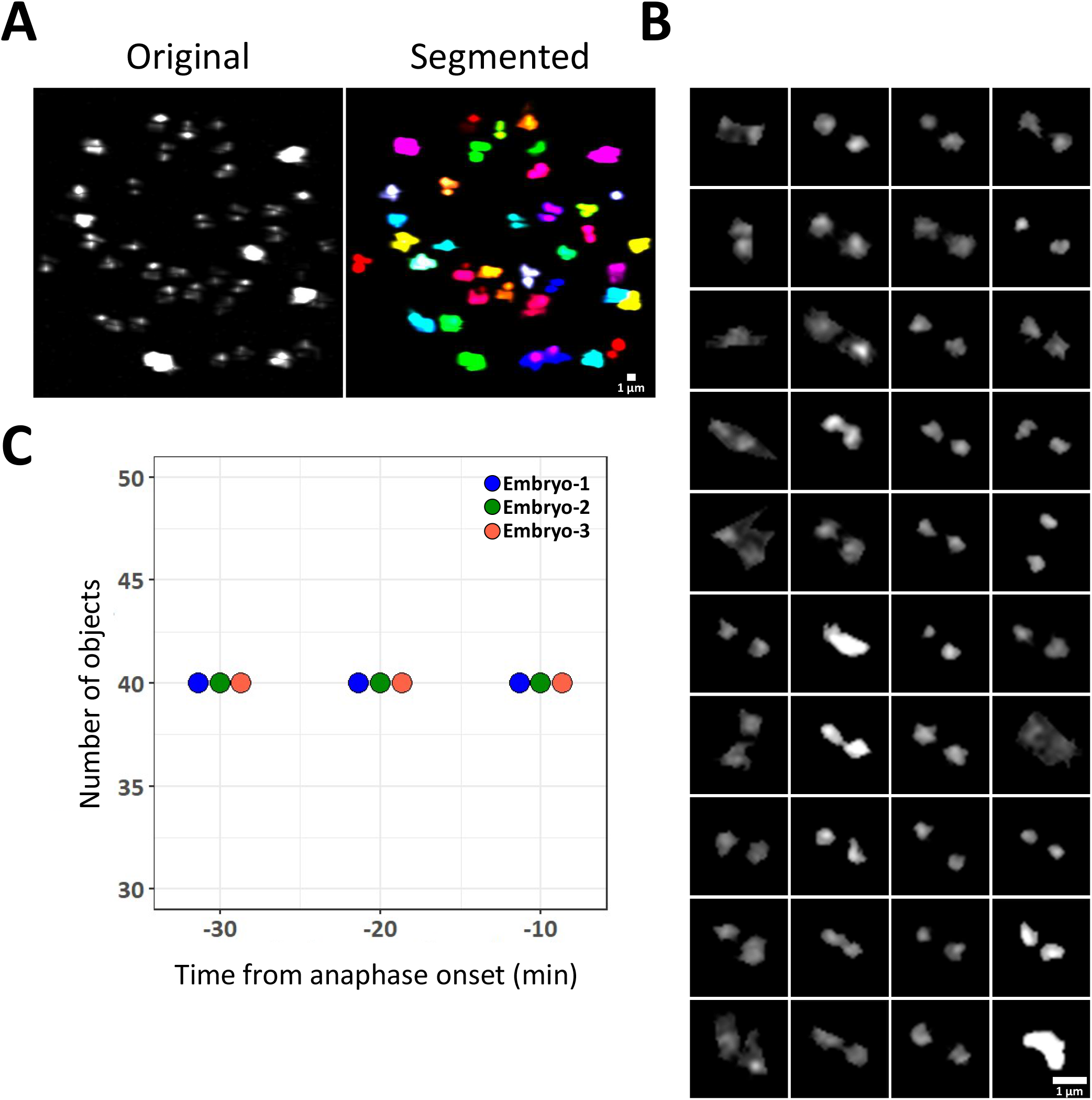
CRISPR-mediated live-cell imaging of the centromere region in the mouse preimplantation embryo. (**A**) Left panel shows the centromere region and right panel shows the segmented image. (**B**) Segmented 40 centromere pairs. (**C**) The number of pairs of dot signals was measured at three points prior to the start of anaphase (t = 0). See also Supplementary Figures 3 and 4.

## Discussion

The objective of this study was to establish a method to count the number of chromosomes in the one-cell zygote without the need for cell collection and fixation. We continuously observed the first cleavage of fertilized mouse eggs using an OPRA-type disk confocal super-resolution microscope and identified the observation conditions under which full-term embryos could be obtained. Depending on the image quality and timing, we succeeded in counting M-phase chromosomes using these conditions. Moreover, the CRISPR-mediated live-cell imaging of the minor-satellite region dramatically improved the accuracy of chromosome counting.

### Implications of phototoxicity assessment and full-term development

When observing a thick sample such as an oocyte, the image becomes darker and the resolution in the Z-direction becomes lower than that of a thinner sample, such as most somatic cells. Although these challenges can be overcome by increasing the laser intensity or the number of images obtained in the Z-axis direction, such measures tend to decrease the embryo viability. Moreover, the effect of super-resolution imaging on the viability of preimplantation embryos has not been reported in the few prior studies that utilized such systems (e.g., Airyscan, SIM, and STED) to observe mouse embryos (***Zielinska et al., 2019***). In the present study, because observations using the SoRa system evinced faster fading of H2B-mCherry, which indicated higher phototoxicity than that in conventional systems (***Fig. 2A, B***), we searched for observation conditions that did not affect the prognosis of the embryo. ***Squirrell et al. 1999*** reported that embryo phototoxicity could be evaluated by determining the rate of full-term development following in-utero transplantation of hamster embryos observed using two-photon excitation microscopy. This method is convincing since it can confirm that the observed phenomenon is not caused by dying cells. In the same manner, phototoxicity consequent to observation using a wide-field microscope (***Yamagata et al., 2005***) and a disk confocal system (***Yamagata et al., 2009a***) has been evaluated. In these studies, EGFP-α-tubulin and histone H2B-mRFP1 were injected into fertilized eggs and observed in two colors (excitation wavelengths: 488 nm/561 nm). In comparison, in the present study, using super-resolution microscopy, we found that 2-cell embryos could not grow into blastocysts when observed with an excitation wavelength of 488 nm (***Table. 1***). Conversely, following observation at an excitation wavelength of 561 nm, the cells successfully developed into blastocysts and full-term embryos (***Table. 2***). These findings suggest that the phenomenon observed at the excitation wavelength of 561 nm did not reflect the abnormal behavior of cells dying from phototoxicity. Furthermore, as we identified conditions under which embryo development proceeded to full-term following super-resolution observation, it should be possible in future experiments to associate phenomena observed using the SoRa system with the prognosis of embryo development and the acquisition of offspring.

### Possibilities afforded by super-resolution live-cell imaging of the first mitosis or chromatin

In the present study, we observed the first division of the mouse embryo (***Fig. 3A, B***). Failure of chromosomal segregation during the first mitosis constitutes a major factor in the termination of embryonic development (***Yamagata et al., 2009b***, ***Mashiko et al., 2020***); thus, detailed observation of the first mitosis using super-resolution live-cell imaging will allow for the prediction of developmental failure. We have previously identified a mitotic pattern that causes aneuploidy using a combination of next generation sequencing and low-magnification live-cell imaging (***Mashiko et al., 2020***). By repeatedly associating super-resolution observation of the first mitosis with genomic testing by next generation sequencing methods, it may be possible to clarify the behavior of chromosome segregation, which is prone to producing chromosome abnormalities.

Notably, analysis of the image data obtained in this study revealed that the SoRa system provided sufficiently high resolution to observe chromatin density at the 2-cell stage (***Fig. 4***), which suggests that this system offers considerable promise as a research tool. Since we have established a low-invasive super-resolution observation method, it has become possible to link the images with chromatin information in combination with conventional methods for examining chromatin. For example, the combination of a method for detailed observation of chromatin dynamics that does not affect embryogenesis (***Fig. 4***) with the profiles of chromatin fold structure as observed by chromosome conformation capture (3C)/Hi-C (***Ke et al., 2017***) and transcription as determined by live-imaging (***Bertrand et al., 1998; Park et al., 2014***) will be expected to accelerate chromatin research.

### Application of chromosome counting to zootechnical science and ART

Counting the chromosomes of mouse zygotes from the obtained super-resolution imaging data (***Fig. 5A, B***) successfully allowed the detection of 40 objects (i.e., 40 pairs of sister chromatids) with a risk of miscounting (***Fig. 5C***). The concept that chromosome counts could be obtained using super-resolution microscopy had previously been tested on U2OS cells (***Gao et al., 2012***); however, the effects on cell viability or risk of miscounting were not reported. In the present study, by further applying the CRISPR-mediated live-FISH technique to our super-resolution live-cell imaging system (***Fig. 6***), we obtained 40-pairs (mice: ***Fig. 7***) of dot signals without measurement error over time.

This suggests that if the chromosomes of livestock/human fertilized eggs could be segmented in the same manner as mouse embryos and the number of chromosomes can be measured, the risk of transplanting abnormal ploidy embryos may be reduced. Since the nuclear envelope breakdown (NEBD) timing is different for each embryo, live-cell imaging is useful to find the different timings more easily. Moreover, counting at multiple time points prevents overlooking or overcounting the number of chromosomes.

Although chromosome counting has been attempted by observing the centromere and kinetochore (***Chiang and Lampson, 2013***), to our knowledge the number of chromosomes has not yet been counted in living cells. Notably, the human alpha-satellite sequence (***Willard, 1985***) is homologous to the mouse minor satellite sequence used in the present study; cattle are also recognized to carry centromere satDNA (***Macaya et al., 1978; Modi et al., 1993; Modi et al., 1996***). Thus, human and bovine embryo chromosomes could also be counted using a gRNA that targets the alpha-satellite. Taken together, our methodology presented in this report, the combination of SoRa system and live-FISH technology, has the potential for use in many animal species and shows considerable promise for widespread application in ART and breeding livestock animals.

## Material and Methods

### Animals

This study conformed to the requirements of the Guide for the Care and Use of Laboratory Animals. All animal experiments were approved by the Animal Care and Use Committee at the Research Institute for Kindai University (permit number: KABT-28-001). ICR strain mice (15-week-old) were obtained from Japan SLC, Inc. (Shizuoka, Japan), and were bred under a specific pathogen-free environment. Room conditions were standardized with temperature maintained at 23 °C, relative humidity at 50%, and 12 h/dark: 12 h light cycle. Animals had free access to water and commercial food pellets. Mice used for the experiments were euthanized by cervical dislocation.

### *In vitro* fertilization

To obtain the unfertilized egg, superovulation was induced in female ICR mice (15 weeks old) by intraperitoneal injections of 10 IU pregnant mare serum gonadotropin (ASKA Animal Health Co., Ltd., Tokyo, Japan) and 10 IU human chorionic gonadotropin (hCG) (ASKA Animal Health) at 48 h intervals. Cumulus-intact oocytes were recovered at euthanasia 15–17 h following hCG injection; these oocytes were arrested at metaphase II of the meiotic cell cycle. Spermatozoa were collected from the cauda epididymis of male ICR mice (11 –16 weeks old) in 0.2 mL droplets of TYH medium (***Toyoda et al., 1971***) and capacitated by incubation for 1.5 h at 37 °C under 6% CO_2_ in humidified air. Cumulus-intact oocytes were collected in 0.2 mL of TYH medium and inseminated with spermatozoa (final concentration 75 sperm/μL). Upon insemination, oocytes arrested at metaphase II restarted the meiotic cell cycle and proceeded to interphase in approximately 3 h. After 1.5 h incubation at 37 °C under 6% CO_2_ in humidified air, the cumulus cells were dispersed by brief treatment with hyaluronidase (Type I-S, 120-300 units/mL; Sigma-Aldrich, St Louis, MO, USA). A total of 30 female and 6 male mice were used, with three experimental replications (female 10). The obtained fertilized eggs were frozen after mRNA injection. In this process, the eggs of each group used in the experiment were randomized, which was expected to mask individual differences in mice. Frozen eggs were used as follows: 20 for the examination of fading, 57 for the examination of developmental capacity, 45 for the examination of birth rate, 10 for observation of 1st mitosis, 10 for W1 and SoRa comparison, 10 for chromosome counting using H2B-mCherry, 10 for chromosome counting using CRISPR-mediated live-FISH, and 20 for measuring the distance between sister chromosomes.

### Preparation of mRNAs to express proteins of interest

After linearization of the template plasmids (pcDNA3.1-polyA83: ***Yamagata et al., 2005***, pTALYM3 (Addgene #47874): ***Miyanari et al., 2013***) at the XhoI (histone H2B-mCherry, histone H2B-EGFP) and ApaI (pTALYM3) site, mRNA was synthesized using RiboMAX™ Large Scale RNA Production Systems-T7 (Promega, Madison, WI, USA). For efficient translation of the fusion proteins in embryos, the 5’-end of each mRNA was capped using the Ribo m7G Cap Analog (Promega), according to the manufacturer’s protocol. To circumvent the integration of template DNA into the embryonic genome, the reaction mixtures for in vitro transcription were treated with RQ-1 RNase-free DNase I (Promega). Synthesized mRNAs were treated with phenol-chloroform to remove protein components. The mRNAs were further purified by filtration using MicroSpin™ S-200 HR columns (GE Healthcare, Chicago, IL, USA) to remove unreacted substrates (RNA reaction intermediates) and then stored at −80 °C until use.

### Preparation of the dCas/gRNA complex

The target sequence of minor satellites of mouse was 5’-ACACTGAAAAACACATTCGT-3’ (***Anton et al., 2014***) crRNA and tracrRNA-ATTO550/ATTO647N (Integrated DNA Technologies, Redwood City, CA) hybridized using a T100 thermal cycler (Bio-Rad Laboratories, Hercules, CA, USA) (94 °C: 5 min, 60 °C: 5 min) were mixed with dCas protein (Integrated DNA Technologies, Redwood City, CA). The final concentrations of gRNA and dCas for mice embryos were 20 and 200 ng/μL, respectively.

### Microinjection of mRNA or dCas/gRNA complex

Probe injection into fertilized eggs was performed as described previously (***Yamagata et al., 2005***). Briefly, mRNAs were diluted to 10 ng/μL using ultrapure water (Thermo Fisher Scientific Barnstead Smart2Pure; Waltham, MA, USA) and an aliquot was placed in a micromanipulation chamber. Fertilized eggs (approximately 4–6 h following insemination) were transferred to HEPES-buffered Chatot-Ziomek-Bavister (CZB) (***Chatot et al., 1989***) medium in the chamber and injected with mRNA using a piezo-driven manipulator with a narrow glass pipette (1 μm diameter). Once the mRNA solution had been aspirated into the pipette, piezo pulses were applied to the fertilized eggs to break the zona pellucida and plasma membrane. A few picoliters of the solution were introduced into the cytoplasm and the pipette was removed gently. The mRNA-injected fertilized eggs were incubated at 37 °C under 6% CO_2_ in air for at least 2 h prior to injection to allow time for protein production. After these procedures, the embryos were frozen.

### Imaging

The motor-control (Mac5000; Ludl Electronic Products, Hawthorne, NY, USA) and piezo control (P-725xDD, PIFOC high dynamics piezo scanner) methods were used to control the Z-axis direction. However, as the images obtained using the piezo-control method were blurred, we adopted the motor-control method for subsequent experiments. The fertilized eggs were transferred to 5 μL droplets of potassium simplex optimization medium with amino acids (KSOMaa) containing 0.00025% polyvinyl alcohol (P8136-250G; Sigma-Aldrich) and 100 μM ethylenediaminetetraacetic acid on a glass-bottomed dish and placed in an incubation chamber (Tokai Hit, Shizuoka, Japan) set at 37°C on the microscope stage. A gas mixture of 5% O_2_, 6% CO_2_, and 89% N_2_ was introduced into the chamber (138 mL/min). An inverted IX-73 microscope (Olympus, Tokyo, Japan) was used with an attached Nipkow disk confocal microscope (CSU-W1 SoRa; Yokogawa Electric, Tokyo, Japan), a Prime95B scientific complementary metal oxide semiconductor (sCMOS) camera (Teledyne Photometrics, Tuscon, AZ, USA), and z-motor (Mac5000; Ludl Electronic Products, Hawthorne, NY, USA). As our imaging device contained an attached auto x-y stage (Sigma Koki, Tokyo, Japan), multiple embryos could be monitored simultaneously. The SoRa system can be switched to the traditional nipkow disk without micro lenses (CSU-W1 mode), allowing parallel observation of the same sample. A set of imaging systems was placed in a dark room with the room temperature maintained at 30 °C. Device control was performed using μ-Manager microscopy software (https://micro-manager.org). When observing in the Z-direction, 101 images were taken at 0.5 μm intervals, 25 μm above and below the equatorial plane of the fertilized egg (total 50 μm). The observed embryos were subsequently moved to an incubator to examine their developmental potential. The power of laser emitted from tip of the objective was measured by optical power meter (TB200; Yokogawa Electric, Tokyo, Japan)

### Image analysis

The 2D images were constructed using MetaMorph software ver. 7.7.10 (Molecular Devices, San Jose, CA, USA), μ-Manager microscopy software, and the ImageJ/Fiji image analysis platform (https://imagej.net/Fiji). The measurements of fluorescence intensity were performed by ImageJ/Fiji. The time constant of the SoRa system was obtained by measuring the time until H2B-mCherry decayed to 36.8% (derived from the definition of the time constant in exponential decay and from the reciprocal of the Napier number), and the time constant of the W1 system was derived from the time until H2B-mCherry decayed to 36.8% after fitting the exponential curve. Centromere images obtained using the W1 and SoRa systems were binarized using the yen-method, denoised using median-filter, and the region of interest (ROI) was detected using Icy (http://icy.bioimageanalysis.org/).

### Chromosome counting/dot signal counting

The FIJI distribution of ImageJ (***Schindelin et al., 2012***) was used for image processing. Deconvolved images were acquired using the Tikhonov regularization algorithm in the DeconvolutionLab2 plugin (***Sage et al., 2017***) with a theoretical point spread function calculated using the Diffraction PSF 3D plugin. Following deconvolution, these images were applied to the Top Hat filter in the MorphoLibJ plugin (***Legland et al., 2016***) for noise removal and binarized using Otsu’s algorithm. Proximate objects in the image were separated using the Transform Watershed 3D algorithm in the MorphoLibJ plugin. The dot signals of the minor satellite were counted using FIJI and FluoRender (***Wan et al., 2012;* https://www.sci.utah.edu/software**).

### Embryo transfer

Embryo transfer has been described previously (***Yamagata et al., 2009b***). Two-cell mice embryos were transferred into the oviducts of day 0.5 pseudopregnant mice. At 18 days following the transfer, cesarean section was performed.

### Sample-size estimation

The blastocyst formation rate in mice is approximately 70–90% and is expected to decrease to 0% to 20% when problems such as environmental or embryo damage occur. We determined that a total of n = 5.08 embryos were required to test the hypothesis that blastocyst formation rate is reduced by super-resolution observation by assuming power = 0.8, sig.level = 0.05, blastocyst ratio of control group = 0.8, blastocyst ratio of observed group = 0.1, one-sided test and prop-test. In turn, the birth rate of mice is approximately 40–60% and is expected to be reduced to 0–10% owing to embryonic damage. We calculated that n = 11.08 embryos would be required to test the hypothesis that super-resolution observation reduced the birth rate by assuming the power was 0.8, sig.level = 0.05, term development ratio of the control group = 0.5, term ratio of the observed group = 0.05, one-sided test and prop-test. However, the lack of significant difference under these conditions does not necessarily suggest the lack of effect. Thus, in the present study, we confirmed whether any abnormalities in blastocyst formation were present following observation of the embryos under these conditions and considered whether live offspring could be obtained. Power analysis was performed using R.

### Experimental replication

With super-resolution observation, the number of eggs that could be observed in one experiment was approximately 10. Thus, the number of eggs used in one experiment was approximately 5 in each of the two groups (biological replication n = 5/once). Technical replication was N = 1 in the examination of fading, N = 7 for blastocyst formation rate, N = 5 for term ratio, N = 3 for first mitosis, N = 3 in the comparison between conventional W1 and SoRa systems, N = 3 each for chromosome counting by H2B-mCherry mRNA and by CRISPR-mediated live-FISH, and N = 2 for measuring the distance between sister chromosomes. The technical replication of experiments related to qualitative conclusions and calculation of the time constant was set to 1. By setting a control group for each experimental replication, we confirmed that inconsistent observation results were not obtained for each experimental replication control group.

### Statistical analysis

One-sided proportion test was performed using the prop.test function in R. Statistics on independent repeats were not possible due to the small number of embryos that could be observed each time in the case of super-resolution live-cell imaging system, so we performed statistics on the total number of embryos.

### Competing interests

The authors declare no conflict of interest.

## Author contributions

Yu Hatano: Methodology, Validation, Formal analysis, Investigation, Resources, Visualization, manuscript review & editing; Daisuke Mashiko: Data curation, Formal analysis, original manuscript draft preparation; Mikiko Tokoro: Formal analysis, Investigation, Resources; Tatsuma Yao: Formal analysis, Software; Kazuo Yamagata: Conceptualization, Supervision, manuscript review & editing, Funding acquisition.

## Acknowledgements

We would like to thank Editage for English language editing. We would like to thank Takuya Azuma for producing the image of the SoRa W1 switching system.

## Funding

This work was supported by JSPS KAKENHI Grant Numbers JP25712035, JP25116005, JP18H05528, and JP18H02357 to KY. The funders had no role in study design, data collection and interpretation, or the decision to submit the work for publication.

## Additional files

**Movie 1 (Related to Figure 3).** Super-resolution observation of the first mitosis of a mouse embryo. The left panel shows the movie from bright field (BF) imaging while the right shows the movie of the histone H2B-mCherry signal.

**Movie 2 (Related to Figure 5).** Three-dimensional (3D) construction of segmented chromosomes. Rotated 3D images of the histone H2B-mCherry signal (left) and processed image (right) are shown.

**Movie 3 (Related to Figure 7).** Live-cell imaging using minor satellite targeted gRNA. Imaging of the minor satellite region for approximately 17 h from the one-cell to two-cell stage.

**Movie 4 (Related to Figure 7).** Three-dimensional (3D) construction of the segmented centromere region. Rotated 3D images of CRISPR-mediated minor satellite signal (left) and the segmented centromere region (right).

**Movie 5 (Related to Figure 7).** Multi color live-cell imaging of a mouse embryo using histone H2B-mCherry and minor satellite-targeted gRNA. The left panel shows the bright-field movie while the right shows the merged movies of histone H2B-mCherry and CRISPR-mediated minor satellite signals.

## Notes

### Competing Interest Statement

The authors have declared no competing interest.

