## Supplemental figures for "Chromosome counting in the mouse zygote using low-invasive super-resolution live-cell imaging"

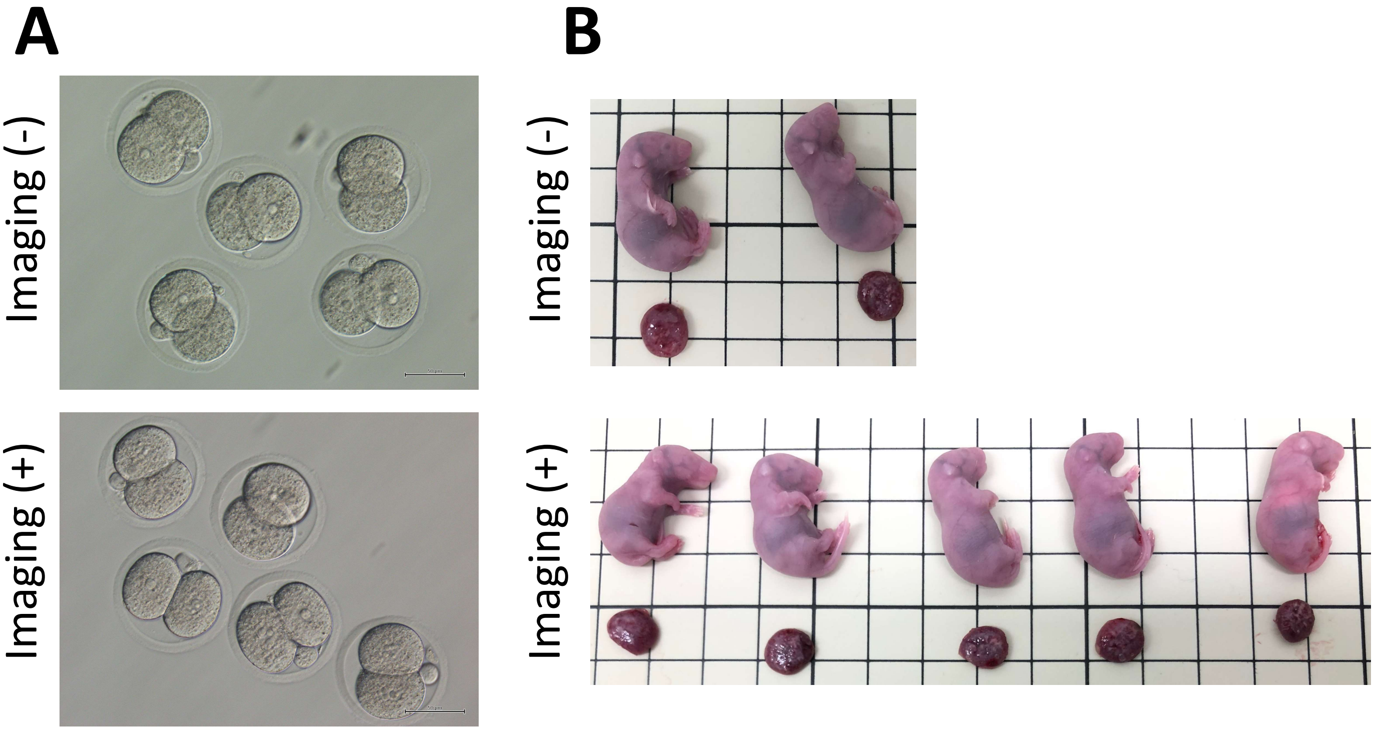


Supplementary Figure 1 (Related to Table 2).

**Supplementary Figure 1 (Related to Table 2).** Two-cell stage embryos prior to transplantation and pups obtained following transplantation. (**A**) Representative image of the 2-cell embryos following observation with an excitation laser at 561 nm, laser power 0.1 mW, and interval 10 min/imaging (−). (**B**) Representative image of the pups obtained following super-resolution imaging. The upper panel shows images of pups derived from imaging (−) and the lower shows an image of pups derived from imaging (+).

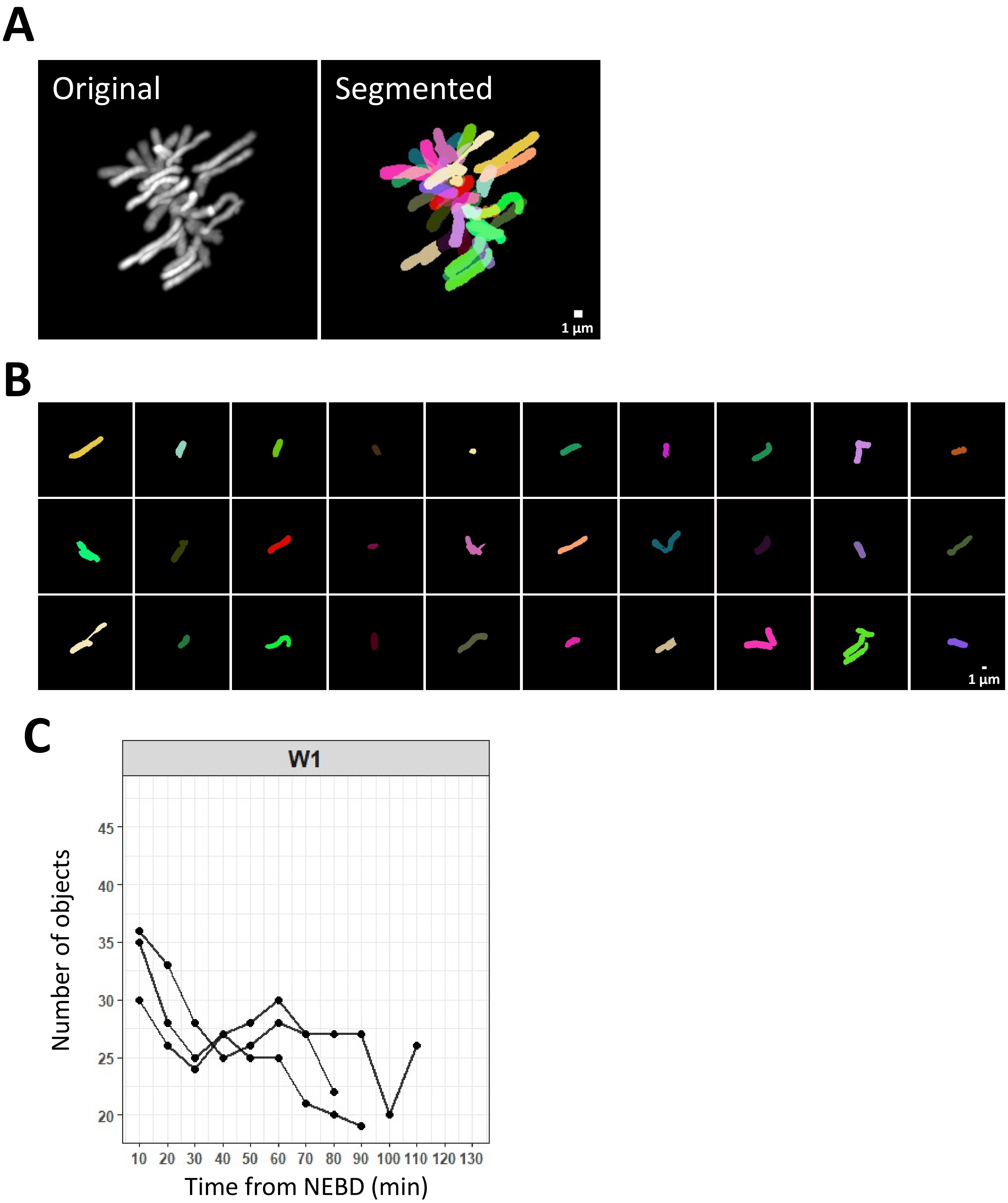


Supplementary Figure 2 (Related to Figure 5).

**Supplementary Figure 2 (Related to Figure 5).** Chromosome counting using the conventional W1 system. (**A**) The left panel shows the histone H2B-mCherry signal. The right image shows the segmented image. (**B**) Each panel shows the recognized object. (**C**) Relationship between the number of recognized objects and time from nuclear envelope breakdown (NEBD).

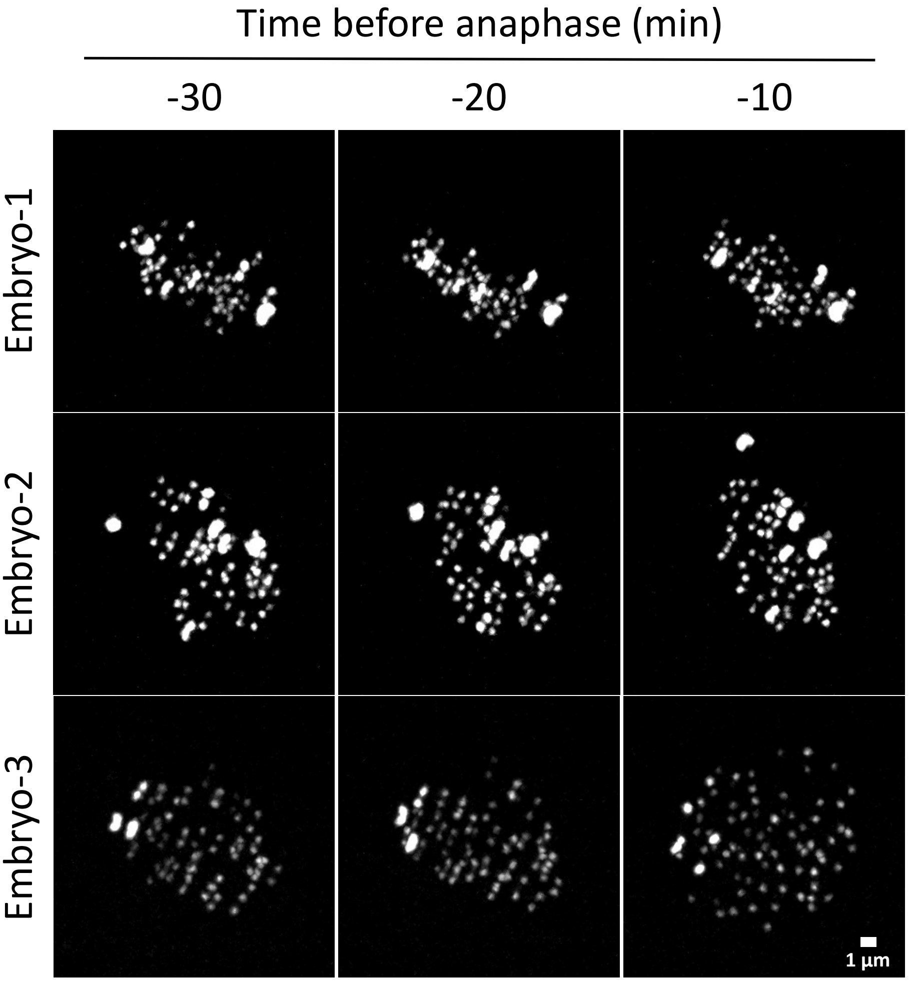


Supplementary Figure 3 (Related to Figure 7).

**Supplementary Figure 3 (Related to Figure 7).** Minor satellite signal of CRISPR-mediated live-cell imaging. Imaging of a minor satellite with the start of anaphase at t = 0. An image of the three time points from each of three embryos is shown.

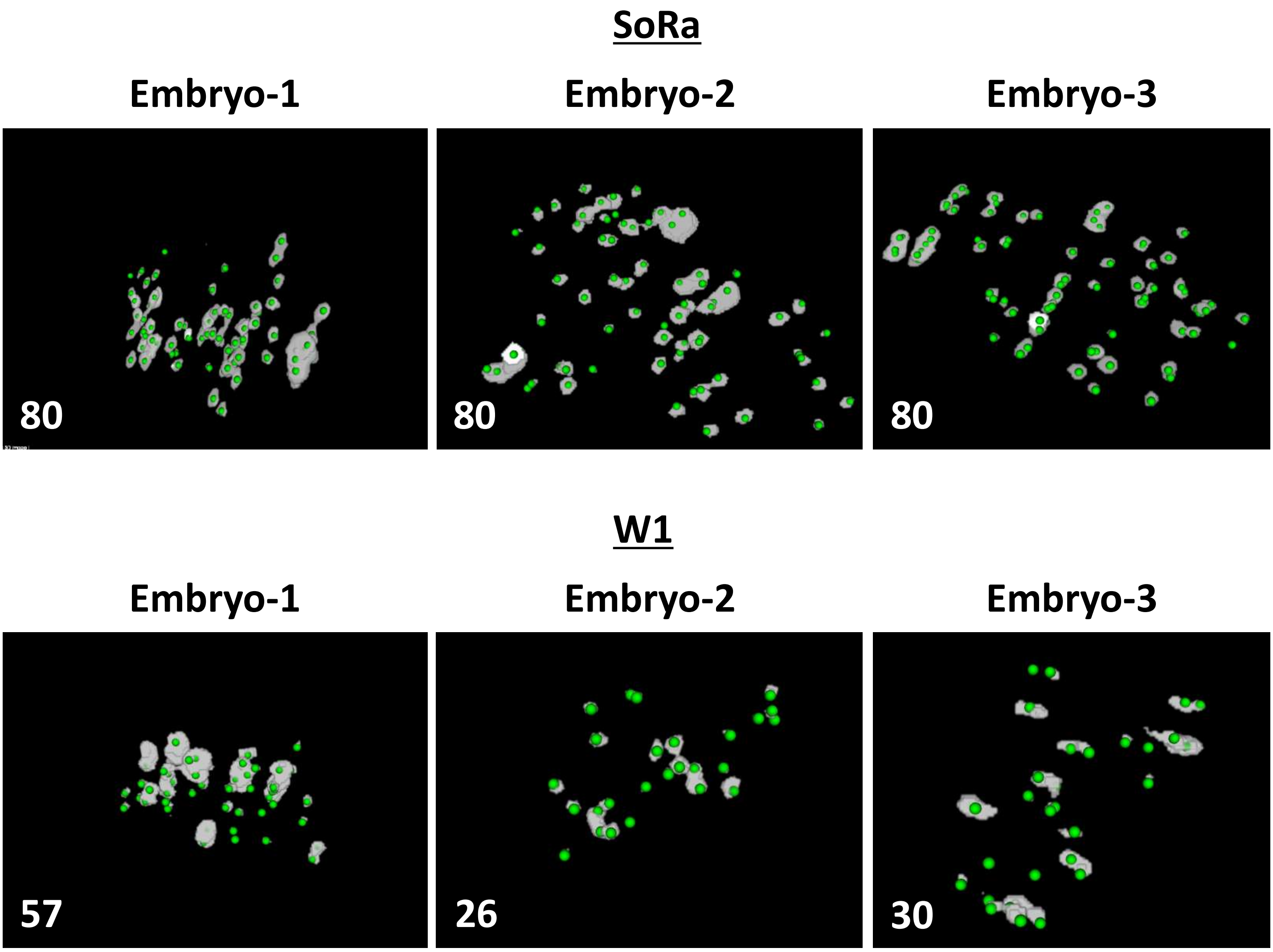


Supplementary Figure 4 (Related to Figure 7).

**Supplementary Figure 4 (Related to Figure 7).** Comparison of detected signal of centromere between SoRa system and W1 system. Upper panels show the binarized images of CRISPR-mediated centromere images obtained using the SoRa system (white) and detected signals (green). Bottom panels show the binarized images of CRISPR-mediated centromere images obtained using the W1 system (white) and detected signals (green). The number in the lower left is the number of dot signals detected.

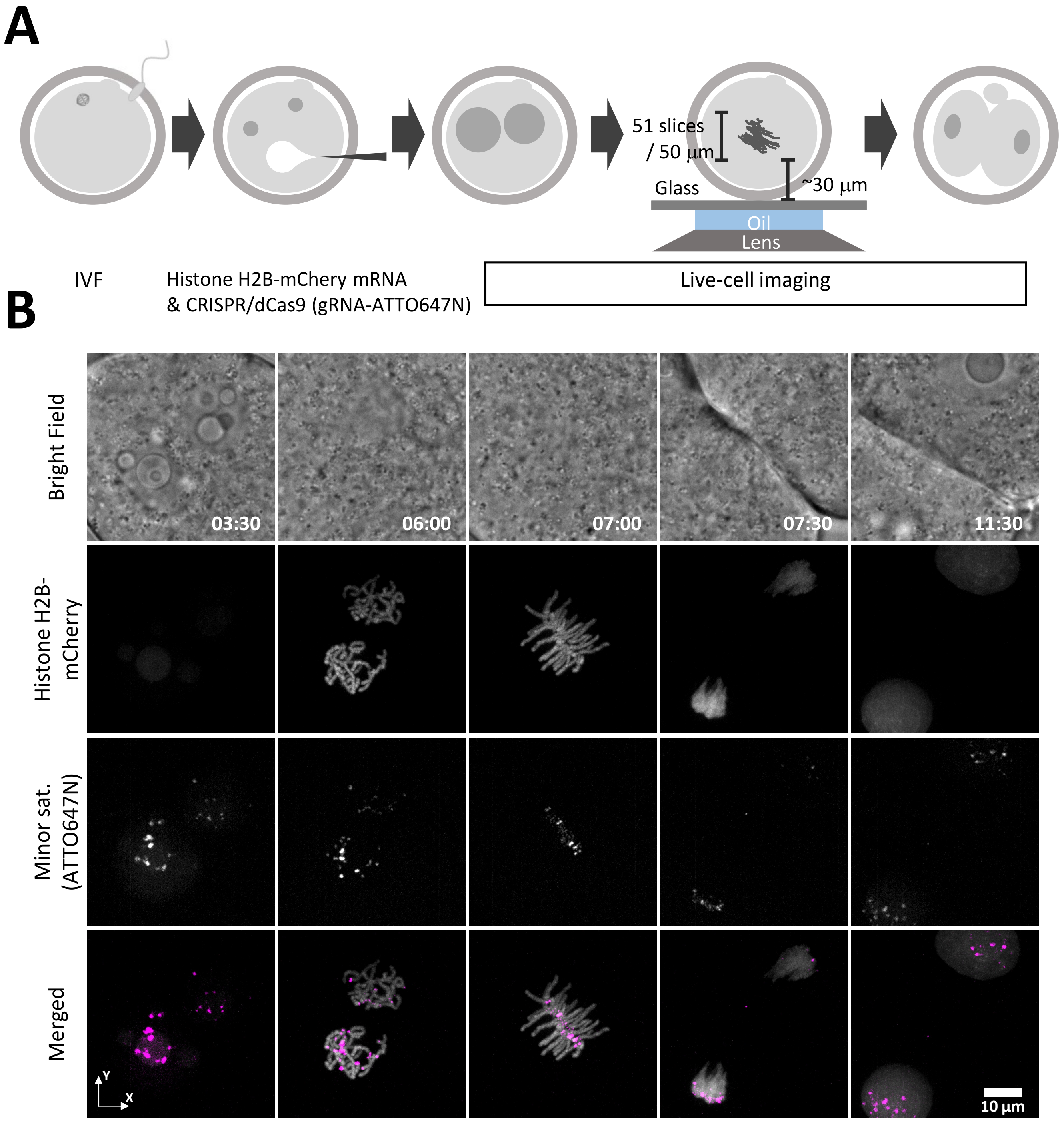


Supplementary Figure 5. (Related to Figure 7).

**Supplementary Figure 5 (Related to Figure 7)** Super-resolution live-cell multi-color imaging of chromosome and Minor satellite. (**A**) Schematic diagram of super-resolution multi color imaging of the first mitosis. (**B**) Super-resolution multi color imaging of the first mitosis in mouse embryos. The upper panels show the bright-field images. Middle panels show the histone H2B-mCherry images of the x–y plane and the panels below show the CRISPR-mediated centromere images of the x–y plane. Bottom panels show the merged images of the x–y plane. See also Supplementary Movie 8.
